# QUANTITATIVE PROTEOMICS OF PLASMA EXTRACELLULAR VESICLES REVEALS A TTR-PLASMINOGEN NETWORK IN ATTR CARDIAC AMYLOIDOSIS

**DOI:** 10.1101/2025.06.28.662124

**Authors:** Amira Zaroui, Damien Habert, Benoit Vallée, Mounira Kharoubi, Soraya Fellahi, Benoit Vingert, Michel Seve, Ilaria Cascone, Vincent Audard, Emmanuel Itti, Hajer Abroud, Arnaut Galat, Thibaud Damy, Sandrine Bourgoin-Voillard, Valérie Molinier-Frenkel

## Abstract

**Background:** Despite recent progress, the prognosis of patients with transthyretin (TTR) cardiac amyloidosis remains poor; this is primarily due to late diagnosis, when irreversible damage has already occurred. Today’s diagnostic work-up still relies on peripheral tissue or a cardiac biopsy, while circulating levels of TTR or other plasma markers have little diagnostic value. Although extracellular vesicles (EVs, as key mediators of intercellular communication) may reflect disease-specific molecular changes, their protein cargo has not yet been explored in the context of TTR amyloidosis (ATTR) cardiomyopathy.

**Objectives:** To characterize the plasma EV proteome in ATTR cardiomyopathy and identify potential biomarkers for pathophysiological pathways, diagnosis, or prognosis.

**Methods:** We performed mass-spectrometry-based, label-free, proteomic profiling of plasma EVs from 65 patients with hypertrophic cardiomyopathy due to TTR amyloidosis (the ATTR+ group, n=41) or non-amyloid cardiac disease (the ATTR- group, n=24). The groups were matched by age and sex.

**Results:** A distinct protein signature comprising 117 deregulated proteins was identified in EVs from ATTR+ patients. The ATTR+ EVs were enriched in proteins associated with vascular homeostasis, coagulation, and inflammation. At least 18 of these proteins formed an interconnected network centered on plasmin/plasminogen. Notably, EV levels of TTR and plasminogen levels were elevated, while the level of alpha2-antiplasmin (plasmin’s primary inhibitor) was low. This imbalance is particularly relevant because plasmin is known to promote amyloidogenesis via TTR cleavage.

**Conclusions:** Our findings provide new insights into the molecular mechanisms underlying ATTR cardiomyopathy and suggest that plasma EV proteins are potential diagnostic or prognostic biomarkers and/or therapeutic targets.

**CONDENSED ABSTRACT:** TTR amyloidosis (ATTR) causes severe cardiac damage, which is often diagnosed late. Through a comparative proteomic analysis of plasma extracellular vesicles (EVs) in patients with ATTR cardiomyopathy vs. patients with other cardiomyopathies, we identified several proteins of relevance to the pathophysiology of ATTR. Our analysis is the first to have highlighted an enrichment of plasmin/plasminogen (known to initiate the amyloidogenic process) and TTR in circulating EVs. Our results might foster the development of (i) diagnostic and prognostic markers for ATTR cardiomyopathy that do not require invasive procedures, and (ii) new therapeutic strategies.

**CENTRAL ILLUSTRATION - GRAPHICAL ABSTRACT:** 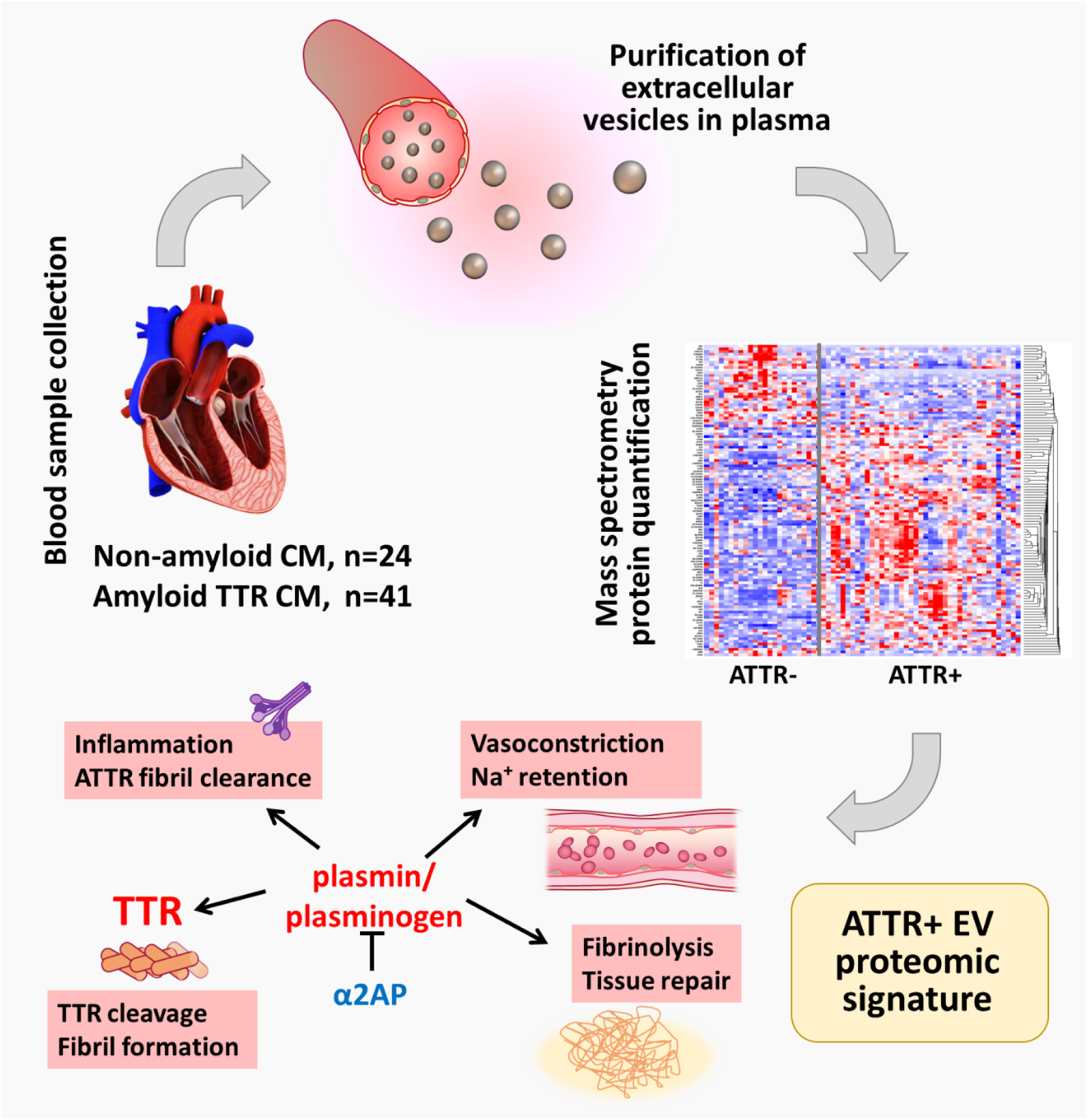

**ETHICAL APPROVAL:** The present analysis was based on blood samples collected as part of a research project entitled "Study of the myocardial microenvironment and toxicity of amyloid proteins in patients with cardiac amyloidosis", which was approved by an institutional review board (CPP Sud-Méditerranée II, Marseille, France; approval references: 2021T2-12/2021-A00950-41 and 2022-A02416-37).

## INTRODUCTION

Amyloidosis encompasses a heterogeneous group of infiltrative diseases characterized by the extracellular deposition of insoluble fibrils. These fibrils are formed by the aggregation of misfolded autologous proteins, the nature of which defines the type of amyloidosis.^1^ The infiltration of organs (such as the heart) by amyloid fibrils affects tissue structure and function. One of the most prevalent forms of cardiac amyloidosis results from the deposition of transthyretin (TTR) either its wild-type or variant form. TTR is a homotetrameric plasma protein that transports thyroxine and vitamin A. However, gene mutations and ageing can limit TTR clearance and/or facilitate dissociation of the tetramer. Accumulation of TTR monomers *in vivo* is followed by their partial unfolding and aggregation to form disease-inducing amyloid fibrils. ^2,3^

TTR amyloidosis (ATTR) cardiomyopathy is a restrictive cardiomyopathy caused by TTR amyloid deposits in the heart. ATTR cardiomyopathy has a poor prognosis, with life expectancy typically ranging from 2 to 6 years after diagnosis. Although ATTR cardiomyopathy was once considered a rare disease, recent advances in diagnostic techniques have revealed that the prevalence has been significantly underestimated.^4,5^ The World Heart Federation acknowledges that improving survival in patients with ATTR cardiomyopathy depends on early diagnosis, appropriate management, a deeper understanding of its pathophysiology, and the development of targeted therapies.

Mass-spectrometry-based proteomic analyses of samples of cardiac tissue has demonstrated its effectiveness in diagnosing ATTR cardiomyopathy.^6,7^ Nevertheless, several studies failed to evidence ATTR-specific protein markers in minimally invasive samples of body fluids, such as blood.^8^ The plasma TTR concentration is not considered to be a relevant diagnostic biomarker *per se*, although the value is often lower in patients than in healthy subjects.^9^

Extracellular vesicles (EVs) are fundamental mediators of intercellular communication; they influence cellular and molecular functions by transporting a variety of biomolecules, including proteins, lipids, and RNAs. Notably, EVs have many prominent physiological and pathological roles and act as key vectors for the transfer of information between cells. As such, EVs modulate complex biological processes (such immune responses, cell proliferation, cell differentiation, and extracellular matrix (ECM) remodeling), depending on their functional interactions. ^10,11^ Thus, a better understanding of the EVs’ functions might lead to many medical applications, such as the identification of new protein biomarkers and the development of delivery systems for the treatment of various diseases ^12,13,14^, including cardiovascular conditions. ^15–18^ Even though the plasma EV protein cargo in ATTR cardiomyopathy might markedly influence disease progression and might have diagnostic value (since TTR is present in EVs), this topic has not been explored previously.^19–23^

At the start of the present study, we hypothesized that cardiac amyloidosis could alter the proteome of circulating EVs. Specifically, we suspected that EVs released by cells stressed by amyloidosis-induced modifications of their microenvironment present specific molecular signatures reflecting the extent and nature of tissue damage. Thus, we conducted a prospective analysis of the proteome of plasma EVs in patients referred to our center for suspected cardiac amyloidosis. We compared patients with a definitive diagnosis of wild-type ATTR or variant form ATTR (ATTRwt and ATTRv, respectively, forming the ATTR+ group) with patients suffering from non-amyloid heart failure (forming the ATTR- group).

## MATERIALS AND METHODS

### 1. Patients and blood sample collection

Patients referred to the French National Reference Center for Cardiac Amyloidosis (at Henri Mondor Hospital, Créteil, France) for suspected ATTR cardiomyopathy and heart failure were enrolled prospectively. The initial clinical suspicion was based on heart failure (with preserved or reduced ejection fraction), imaging features suggestive of cardiac amyloidosis (on echocardiography or MRI), and/or positive diphosphonate scintigraphy with a left ventricular wall thickness ≥12 mm. Patients with grade 1 radiotracer uptake on scintigraphy or monoclonal gammopathy underwent a cardiac biopsy for confirmation of ATTR, in line with the expert consensus.^24^ The following data were recorded for each patient: the personal medical history, ongoing medications, demographic variables, clinical variables (weight, height, heart rate, blood pressure, electrocardiogram, clinical signs of heart failure such as New York Heart Association status), cardiac events, and extracardiac events), and laboratory variables (high-sensitivity cardiac troponin, N-terminal pro-brain natriuretic peptide (NT-proBNP), creatinine, hepatic and renal biomarkers, proteinuria, C-reactive protein (CRP), plasma TTR, complete blood count, coagulation profile, and tests for monoclonal gammopathy). The echocardiographic assessment included cardiac output and left ventricular variables (Table I). Two blood samples were collected on EDTA tubes for each patient at first visit and 1 mL plasma was separated by centrifugation and immediately stored at -80°C for subsequent EV isolation and proteomic analysis.

Patients with a confirmed diagnosis of ATTR cardiomyopathy (the ATTR+ group) were classified as ATTRwt or ATTRv, depending on the status of the *TTR* gene. As mentioned above, patients with non-amyloid non-infiltrative myocardiopathy (the ATTR- group) constituted the control group.

The main exclusion criteria were age under 18 years, a diagnosis of non-ATTR cardiomyopathy or another infiltrative myocardiopathy, treatment with oncolytic chemotherapy or immunosuppressant drugs, liver cirrhosis, renal dialysis, and the lack of social security coverage. All study participants were diagnosed and followed up in our center. After excluding patients based on predefined exclusion criteria, we selected control subjects (ATTR- patients) with non-amyloid non-infiltrative myocardiopathy and who were approximately matched to ATTR+ patients (primarily based on age and sex) in a 2:1 ratio. This matching was intended to minimize potential confounding due to demographic differences between the groups.

### 2. EV isolation and characterization, and the proteomics analysis

To quantify the proteome in EVs, we used a previously described protocol ^25^ that had been modified in accordance with guidelines from the International Society for Extracellular Vesicles (for details, see Supplemental data 1).^26–28^ Briefly, EVs were isolated from 1 mL of plasma by differential ultracentrifugation and then characterized using flow cytometry. The EV proteome was identified and quantified using a differential, label-free, quantitative mass-spectrometry approach. The latter was based on a bottom-up protocol, with trypsin and lys-C digestion and a nano-liquid chromatography mass spectrometry (LC-MS) analysis (NanoAcquity-ESI-SynaptG2-Si, Waters, Milford, MA, USA).^25,29^

The samples’ protein cargos were compared with publicly available EV proteome libraries (ExoCarta July_2015_Release ^21–23^ and/or Vesiclepedia August_2018_Release ^19,20^), using FunRich software (version 3.1.3).^30,31^ Downstream analyses were performed with Gene Ontology (GO), pathway and disease enrichment, all generated by StringApp (version 2.1.1, ^32^ implemented in Cytoscape (version 3.9.1)). ^33^ Characteristics of extracellular vesicles (EVs) isolated from plasma samples were analyzed in ATTR+ and ATTR- patients.

The number of EVs in plasma was assessed by flow cytometry, using Trucount™ beads as previsouly described. ^25^ EVs were first identified by CD9 labeling. Megamix fluorescent beads (Biocytex, La Ferté Saint-Aubin, France) were used to differentiate between particles with diameters of 160, 200/240, 300, 500 and 900 nm. The flow cytometry data were analyzed with FlowJo software (FlowJo, Ashland, OR; version 10.7.1). A Venn diagram was plotted to compare the identified EV proteins with those in the Vesiclepedia and/or ExoCarta EV libraries. A GO enrichment analysis of cellular and extracellular compartments was conducted using StringApp (version 2.1.1) implemented in Cytoscape (version 3.9.

### Proteomic data availability

The study data are provided in the manuscript and the Supplemental Appendix. Proteomic data have been deposited with the ProteomeXchange Consortium (http://proteomecentral.proteomexchange.org) through the MassIVE repository resource (MassIVE MSV000098005), with the dataset identifier PXD064342.

### 3. Statistical Analysis

Patients in the ATTR- group were matched 2:1 by age and sex with those in the ATTR+ group, in order to minimize confounding factors and enhance the robustness of the proteomic analysis. The groups’ clinical characteristics were compared using a t-test, chi-square test, or Mann-Whitney U test, depending on the data distribution.

A quantitative statistical analysis of proteomic data was based on an analysis of variance (Progenesis for the proteomics QI application (Waters) after the signal had been normalized against the internal standard (ENO-1 from yeast). Proteins with a p-value ≤ 0.05, Q-value ≤ 0.05, and an absolute log2 fold-change > 1.5 were considered to be differentially expressed proteins (DEPs) and were included in subsequent analyses. Volcano plots and principal component analyses were performed using R software (version 4.5.0; R Foundation for Statistical Computing, Vienna, Austria) within the RStudio environment (Posit Software, Boston, MA). Dot plots were built with GraphPad Prism software (version 10, GraphPad Software LLC, Boston, MA, USA). Heat maps were constructed using FunRich software (version 3.1.3 a standalone tool for functional annotation and pathway analysis.^30,31^ To highlight relevant biological processes, pathways and diseases, enrichment analyses were performed using a false discovery rate (FDR) threshold ≤ 0.05 in the Benjamini-Hochberg test.

## RESULTS

### 1. Clinical data

Between July 2020 and June 2024, 430 patients with suspected cardiac amyloidosis or a suspected, unrelated myocardiopathy consulted in our department (Supplemental Data 2). One hundred and eleven patients met our criteria for inclusion in the ATTR+ or ATTR- groups, and 65 patients with confirmed ATTR amyloidosis (ATTR+, n=41) or without evidence of ATTR deposition (ATTR−, n=24) were selected using frequency matching based on key demographic and clinical variables, including age, sex, and cardiac phenotype. Exact 2:1 matching was not enforced, in order to maximize the number of available cases while preserving group comparability for relevant covariates. The ATTR+ group comprised 30 ATTRwt cases and 11 ATTRv cases.

Demographic, clinical, laboratory, echocardiographic and medication-related variables in the final cohort were compiled and analyzed statistically (Table 1). Although the ATTR+ or ATTR- groups were well matched for most features, we observed differences in the plasma levels of gamma-glutamyl transferase (GGT) and CRP, left ventricle and septum echocardiographic features, and medications (beta-blockers and drugs targeting the renin-angiotensin-aldosterone system). These differences should be borne in mind when interpreting the study’s findings.

**Table 1.** Baseline characteristics of the study population. Using Fisher’s test for categorical variables and the Mann-Whitney test for continuous variables, the threshold for statistical significance was set to p<0.05. **ATTR:** TTR amyloidosis, **NYHA**: New York Heart Association, **HR**: heart rate, **bpm**: beats per minute, **SBP**: systolic blood pressure, **NT- proBNP**: N-terminal pro B-type natriuretic peptide, **eGFR**: estimated glomerular filtration rate, **AST**: aspartate aminotransferase, **ALT**: alanine aminotransferase, **GGT**: gamma-glutamyl transferase, **CRP**: C-reactive protein, **LVEF**: left ventricular ejection fraction, **IVS**: interventricular septum, **GLS**: global longitudinal strain, **PASP**: pulmonary artery systolic pressure, **ACEI**: angiotensin-converting enzyme inhibitor, **ARB**: angiotensin II receptor blocker.

|  | Whole population<br>(n=65) | ATTR+<br>(n=41) | ATTR-<br>(n=24) | P-value |
| --- | --- | --- | --- | --- |
| **Age, median [IQR]** | 77 [69 - 84] | 82 [77-86] | 70 [54 - 79] | 0.12 |
| **Men, n (%)** | 48 (75) | 30 (73) | 18 (75) | 0.60 |
| **Hereditary, n (%)** |  | 11 (26.8) |  |  |
| **Clinical features** |  |  |  |  |
| NYHA class III-IV, n (%) | 21 (33) | 13 (32.4) | 8 (34) | 0.34 |
| HR (bpm), median [IQR] | 74 [68 - 89] | 71 [67 - 89] | 77 [68 - 91] | 0.88 |
| SBP (mm Hg), median [IQR] | 127.0 [113 - 146] | 129 [115 - 149] | 117 [100 - 143] | 0.56 |
| Neurological involvement, n (%) | 14 (21.8) | 10 (24) | 4 (16) | 0.04 |
| **Biomarkers, median [IQR]** |  |  |  |  |
| Prealbumin (TTR, g/L) | 215 [173 - 267] | 206 [172 - 251] | 233 [193 - 279] | 0.03 |
| NTproBNP (ng/L) | 2362 [1377 - 4757] | 2308 [1331 - 4900] | 2394 [1430 - 4873] | 0.67 |
| Troponin (ng/L) | 62 [36 - 87] | 72 [44 - 84] | 54 [31 - 88] | 0.12 |
| eGFR (mL/min/m<sup>2</sup>) | 55.4 [48.6 - 63.2] | 56.8 [49.5 - 63.1] | 53.8 [48.4 - 60.3] | 0.09 |
| AST (U/L) | 23 [18 - 31] | 21 [17 - 25] | 30 [20 - 56] | 0.07 |
| ALT (U/L) | 30 [24 - 34] | 30 [25 - 33] | 30 [23 - 46] | 0.89 |
| GGT (U/L) | 63 [37 - 124] | 56 [34 - 84] | 81 [40 - 180] | 0.04 |
| CRP (mg/L) | 3.7 [1.6 - 7.9] | 4.2 [1.3 - 9.6] | 3.7 [2.9 - 12.3] | 0.04 |
| Platelet count (G/L) | 206 [170 - 244] | 208 [176 - 246] | 189 [159 - 245] | 0.68 |
| **Echocardiographic features, median [IQR]** |  |  |  |  |
| LVEF (%) | 47 [30 - 55] | 50 [42 - 56] | 38 [28 - 54.7] | 0.02 |
| LVEF <35%, n (%) | 14 (22.5) | 5 (12) | 9 (37) | 0.04 |
| IVS thickness (mm) | 15 [12 - 18] | 17 [15 - 18] | 12 [7 - 14] | 0.03 |
| GLS (% absolute value) | 8.5 [6.2 - 10.7] | 8.4 [6 - 17.5] | 9.5 [8.0 - 13.0] | 0.01 |
| PASP (mmHg) | 34 [28 - 42] | 30 [28 - 45] | 36 [30 - 47] | 0.06 |
| **Medications, n (%)** |  |  |  |  |
| Anticoagulants | 39 (61.8) | 25 (62.5) | 24 (58) | 0.55 |
| Furosemide | 46 (72.5) | 30 (73) | 16 (66) | 0.06 |
| ACEIs/ARBs/anti-aldosterone | 44 (70) | 20 (48) | 24 (100) | 0.01 |
| Betablockers | 20 (31) | 0 | 20(83) | - |
| Amiodarone | 13 (20.6) | 8 (28.6) | 5 (20) | 0.06 |

### 2. The plasma EV protein signature in patients with ATTR cardiomyopathy

Plasma EVs were characterized using flow cytometry and LC-MS (for detailed data, see Supplemental Data 3). The EV concentration and size distribution profile were similar in the ATTR+ and ATTR- groups. In both groups, extracellular vesicles ranging from 160 to 900 nm were detected. ^34^ Most of the EVs were between 160 and 250 nm in diameter, indicating a higher concentration of small extracellular vesicles - small microparticles typically derived from the plasma membrane or large exosomes derived from intracellular compartments ^34^- rather than larger, cell-derived microparticles. A large-scale proteomic analysis of the EVs enabled the quantification of 448 proteins. Most of these proteins were related to EVs, as indicated by an enrichment in the extracellular compartment according to the GO analysis and comparison with the Vesiclepedia^19,20^ and ExoCarta^23^ libraries, confirming the high purity of the patients’ EVs and a greater enrichment in proteins derived from the plasma membrane. Circulating EVs represent a complex, heterogeneous system because they can be derived from any cell or tissue in the body. We also characterized the source of the plasma EVs by mapping the tissue and blood distribution of the total protein content (Supplemental Data 4). Most of the quantified proteins were present in the blood, the heart, and the peripheral nervous system, which is commonly infiltrated by TTR amyloid fibrils.^31^ Thus, plasma EVs are valid markers of the proteins involved in ATTR.

Of the 448 quantified proteins in the ATTR+ vs. ATTR- groups, 117 (26%) were DEPs: 85 were upregulated and 32 were downregulated in ATTR+ patients (Figure 1A, table S1). A principal component analysis based on individual patient levels of the 117 DEPs revealed a clear separation between ATTR+ and ATTR- patients (with the first two dimensions accounting for 21.9% and 12.4% of the total variance, respectively). This finding emphasized the distinct proteomic profiles associated with disease status (Figure 1B); which were also visible on a heatmap (Figure 1C). The dot plots (Figure 1D highlighted four highly significant DEPs in ATTR+ EVs: TTR, plasmin/plasminogen (the analysis cannot differentiate between plasmin and plasminogen), α2-antiplasmin, and C1Q C-chain. Plasmin/plasminogen and α2-antiplasmin potentially regulating amyloid fibril deposition directly (see below), whereas C1Q C-chain was one of the six complement proteins upregulated in ATTR+ EVs.

**Figure 1.**
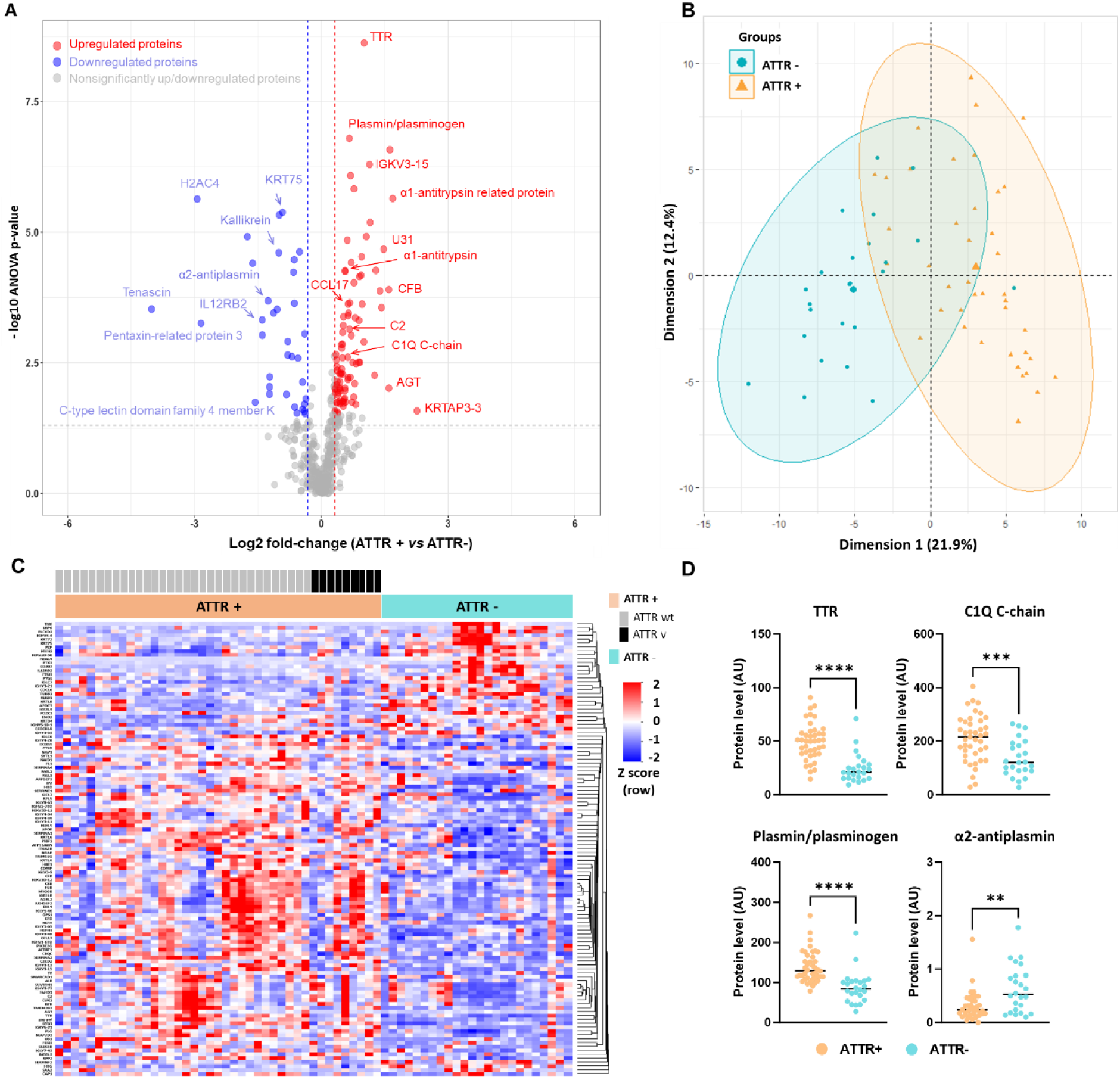
Quantitative proteomics data from plasma EVs in the ATTR+ and ATTR- groups. (**A**) A volcano plot depicting the log2-transformed fold-change and the corresponding -log10 FDR for all quantified proteins. Each dot represents a single protein. The application of an FDR threshold of 0.05 and a fold-change cutoff of 1.25 led to the identification of 117 DEPs. Up- and downregulated DEPs in ATTR+ patients are shown in red and blue, respectively. (**B**) A principal component analysis score plot for the DEPs. Each dot represents an individual’s proteomic profile projected onto the first two principal components (Dimension 1 and Dimension 2). The cluster of patients in the ATTR+ group (in orange) and the cluster of patients in the ATTR- group (in green) patients were clearly separate. (**C**) A heatmap of the 117 DEPs. The color scale represents the range of log2-transformed values, with blue indicating low expression and red indicating high expression. (**D**) Individual raw patient data for levels of TTR, plasmin/plasminogen (the analysis cannot differentiate between plasmin and plasminogen) and α2- antiplasmin (alternate name: SERPINF2). AU: arbitrary units. **** P-value <0.0001, *** P-value <0.001, ** P- value <0.01.

### 3. Gene ontology, pathway and disease enrichment analyses

We next performed GO and pathway enrichment analyses, in order to identify the biological functions and processes that were most strongly associated with the DEPs. The analysis of three independent databases (GO Biological Processes, Reactome Pathways, and WikiPathways) confirmed the major importance of complement activation in ATTR+ EVs (Figure 2 and Table S2). Several other biological processes (mainly related to ECM remodeling, wound healing, coagulation, inflammation, and the humoral immune response) were significantly enriched among the DEPs. The disease enrichment analysis showed that proteins upregulated in ATTR+ EVs could be related to several affections; the most relevant for cardiac amyloidosis being “familial visceral amyloidosis” (Disease Ontology Identifier (DOID):005063), “coronary artery disease” (DOID:3393) and “thrombosis” (DOID:0060903). Familial visceral amyloidosis is a rare, inherited form of amyloidosis caused by mutations affecting various genes, including *TTR*. Overall, the pathways identified through these unsupervised, hypothesis-free analyses highlighted the importance of complement activation, deregulation of the coagulation cascade, and ECM remodeling.

**Figure 2.**
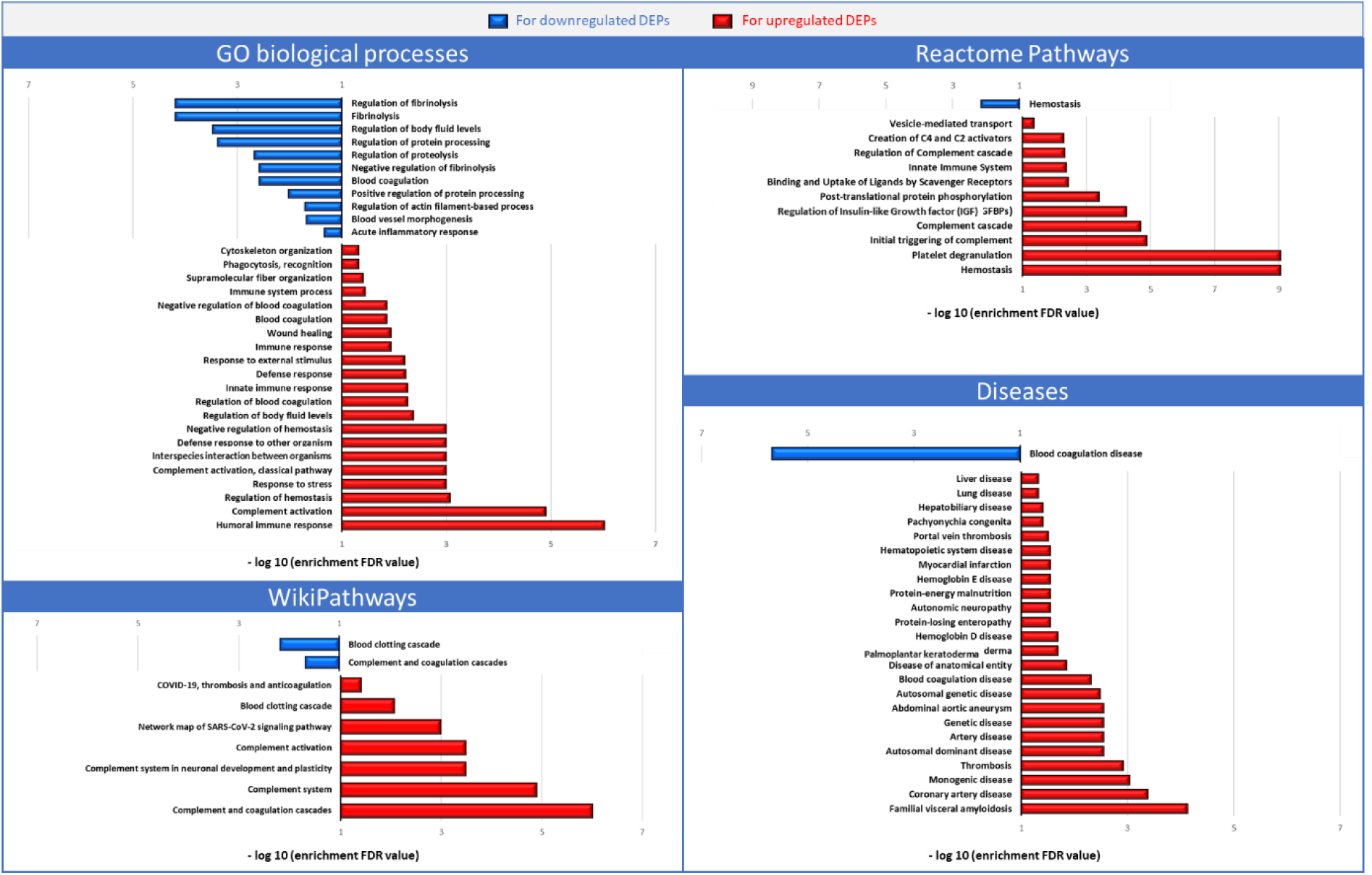
Enrichment analyses of DEPs, using publicly available databases for biological processes and diseases. Analyses were performed using StringApp (version 2.1.1) implemented in Cytoscape (version 3.9.1).

### 4. Plasmin/plasminogen is at the center of the ATTR+ protein signature

A closer examination of the DEPs revealed that 18 belonged to four molecular cascades: the complement system, the intrinsic (or contact) coagulation pathway, and the kallikrein-kinin and renin-angiotensin systems (supplemental data 5). Several proteins from these cascades are functionally interconnected through activation and inhibition mechanisms, which frequently involve protease activity that generates bioactive peptides or enzymes from precursor proteins (see Figure 3, which shows DEPs enriched (or not) in EVs (in red and blue, respectively).

**Figure 3.**
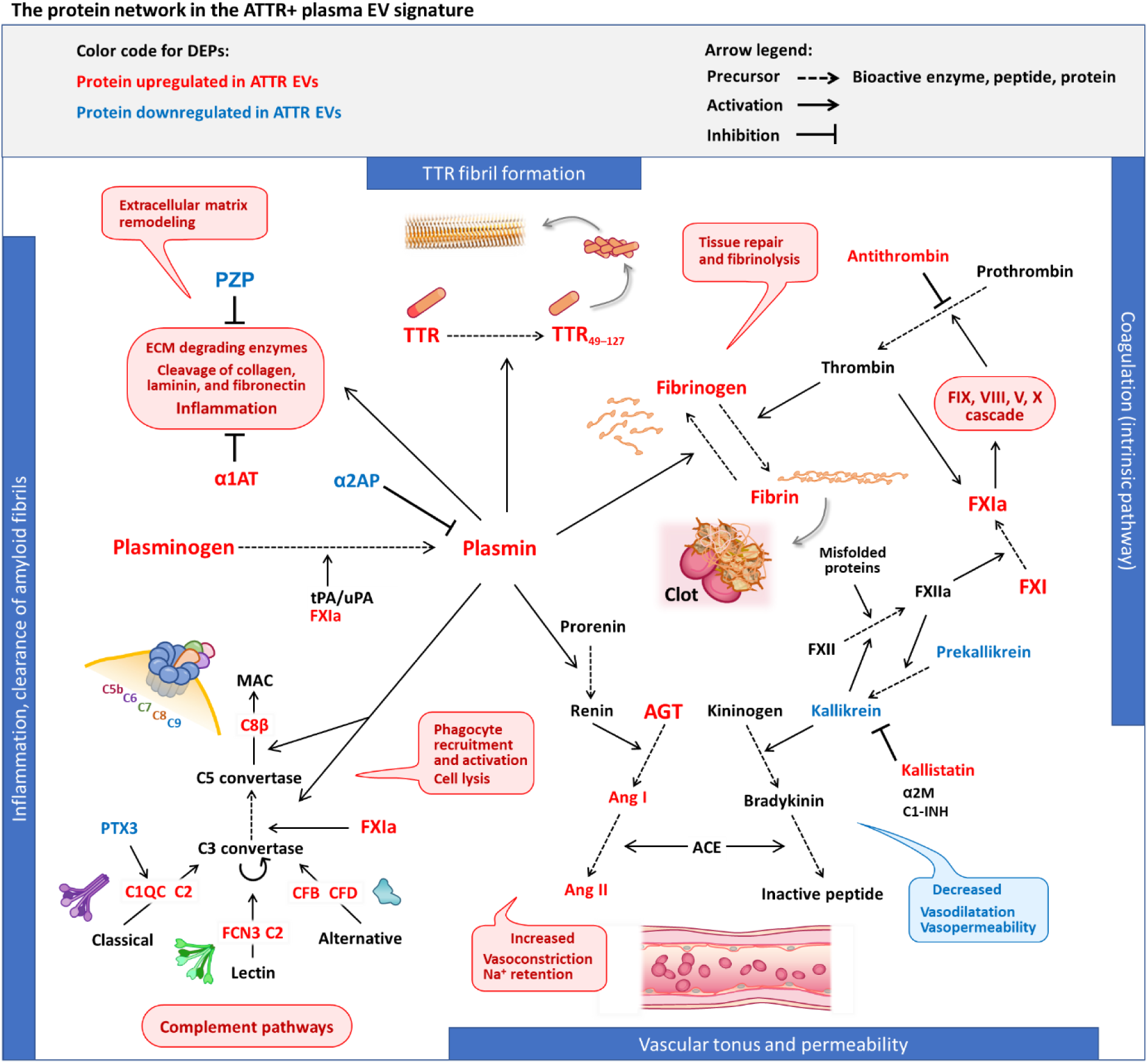
The protein network underlying the ATTR+ plasma EVs’ proteomic signature. The diagram highlights the main relationships between the various proteins involved in coagulation and tissue repair (the intrinsic pathway), inflammation (the complement system), and vascular homeostasis (the renin−angiotensin and kallikrein−kinin systems). The network is centered on plasmin, the activity of which also regulates the formation of TTR amyloid fibrils, ECM remodeling, inflammation and complement factor C5 activation.

At the center of these pathways is the precursor plasminogen and its cleaved product plasmin; like other protein/peptide pairs, a proteomic analysis cannot distinguish between the two. The levels of plasmin/plasminogen were significantly elevated in ATTR+ EVs (ATTR+/ATTR- ratio = 1.58; *p* < 0.00001). The many key roles of this plasma protein (including interactions with protein targets and cell surface receptors, and the serine protease activity of the cleaved form plasmin) are well documented in the literature.^35^ Interestingly, in the context of the ATTR+ signature, plasmin/plasminogen interacts with complement components, inflammatory cells, the ECM, ECM-degrading enzymes, the renin-angiotensin system, and (through its well-characterized fibrinolytic activity) the coagulation cascade. Even more interestingly, it has been shown that plasmin-mediated TTR cleavage liberates a truncated monomer (TTR_49-127_) capable of initiating the formation of amyloid fibrils.^36,37^ We also found that α2-antiplasmin (the main inhibitor of plasmin activity) was strongly deeply downregulated in ATTR+ EVs (ratio ATTR+/ATTR- = 0.48; *p* = 0.00198). Hence, plasmin activity is probably enhanced in ATTR+ EVs, which might lead to TTR cleavage and greater amyloid deposition. Most importantly, TTR was very significantly upregulated in ATTR+ EVs (ratio ATTR+/ATTR- = 2.02; *p* < 0.00001). In contrast, the ATTR+ vs. ATTR- patients did not differ significantly with regard to the plasma TTR concentration (Table 1).

## DISCUSSION

The present study is the first to have explored the proteome of circulating EVs from patients with ATTR cardiomyopathy. We identified a specific ATTR+ signature by comparison with the proteome of circulating EVs from patients with non-amyloid hypertrophic cardiac diseases (i.e. the actual differential diagnosis for ATTR+ in clinical practice). EVs serve as a means of intercellular communication and are likely to offer valuable insights into the processes at work in amyloidosis, independently of total circulating proteins.

The ATTR+ signature comprised 117 DEPs, including TTR and 18 proteins directly involved in four highly intricate molecular systems: the complement system, the renin-angiotensin system, the kallikrein-kinin system, and the coagulation cascade (Figure 3). Several of these proteins may be produced by (i) cells in organs affected by amyloidogenesis and/or (ii) innate immune cells (such as monocytes and macrophages) and activated endothelial cells involved in the inflammatory response. Some of the DEPs have been detected previously in the proteomic signatures of ATTR heart biopsies (apolipoprotein E and TTR) and carpal tunnel tissue (complement proteins).^38,39^ Based on our unsupervised enrichment analyses, similar biological pathways were found to be prominent in EVs and cardiac tissue from ATTR patients. The ATTR+ signature appears to have links to cardiac diseases and amyloidosis. Some of the key proteins in the ATTR+ signature are discussed below.

### TTR

This protein was significantly enriched in ATTR+ EVs, whereas the ATTR+ patients’ plasma concentration was low (as observed previously).^9^ It has been argued that a low plasma TTR is due to rapid capture by amyloid aggregates in target organs.^9^ A direct comparison between plasma TTR and plasma EV TTR levels was limited by our experimental techniques: mass spectrometry (which identifies TTR peptide fragments) and a TTR immunoassay (using TTR-binding antibodies) are both highly specific but cannot distinguish between the native tetramer and dissociated monomers. The lack of an elevated plasma TTR concentration in ATTR+ patients probably reflected the EVs’ cellular origin (the proteome of which tends to reflect tissue-level processes), rather than technical limitations in detection.^40,41^ For instance, EVs derived from mononuclear phagocytes (such as circulating monocytes or tissue macrophages) might be enriched in TTR following the phagocytosis of amyloid deposits - a process potentially facilitated by complement activation.

### Complement system

Along with immunoglobulins (activators of the classical pathway), proteins from the complement cascade were among the most abundant in plasma EVs (as reported previously)^42^. The complement system is essential for innate immunity and inflammation and can lead to cell lysis through formation of the membrane attack complex. Six complement components were specifically enriched in ATTR+ EVs, including complement factor D (CFD). CFD is mainly produced by adipocytes and mononuclear phagocytes (but not by the liver), circulates in an active form, and is highly substrate-specific.^43^ Importantly, CFD is a rate-limiting enzyme with a key role in amplifying C3 convertase activity. Thus, our results suggest that complement cascade was fully activated and might have been amplified in the final steps by plasmin and factor FXI (Figure 3).^44,45^ Our results are in line with previous reports of irreversible ultrastructural damage and complement activation in cardiac tissue, carpal tunnel tissue, and subcutaneous abdominal fat tissue from ATTR patients.^38,39,46^

Conversely, complement activation might facilitate (i) the elimination of cell debris and immune complexes^47^, and (ii) the clearance of amyloid deposits through immunoglobulin-mediated or direct binding of C1Q to amyloid fibrils.^48^

Pentraxin-3 was strongly downregulated in ATTR+ EVs and was much less abundant than complement components. This might have been due to the chronic nature of the inflammation associated with amyloidosis, given that pentraxin-3 is an acute-phase inflammatory protein.

### Plasmin/plasminogen and α2-antiplasmin

Plasminogen and/or plasmin were enhanced in ATTR+ EVs. We suggest that plasmin activity is elevated in ATTR+ EVs (due to a greater level of FXI and a lower level of α2-antiplasmin) and that plasmin/plasminogen has a central role in connecting the pathways involved in the pathogenesis of amyloidosis. Although plasmin is well known for its fibrinolytic role, this serine protease has a broad substrate specificity and can therefore participate in extracellular matrix remodeling, activation of the renin−angiotensin system, and (as mentioned above) complement activation.^47,49^ Moreover, plasminogen can bind to dead cells and misfolded proteins (including amyloid fibrils), and plasminogen receptors have notably been identified at the surface of immune cells and endothelial cells.^35^

Most importantly, plasmin can cleave TTR *in vitro* between residues 48 and 49 to liberate a truncated monomer (TTR_49-127_) that destabilizes the protein and promotes aggregation.^36^ This was confirmed for a mutant human TTR, using a transgenic mouse model of cardiac amyloidosis .^37^ Furthermore, deletion of the *Serpinf2* gene (encoding α2-antiplasmin) in this model accelerated the development of ATTR cardiomyopathy.^37^ Thus, upregulation of plasmin/plasminogen and TTR and downregulation of α2-antiplasmin in ATTR+ EVs may reflect both active amyloidogenesis and an inflammatory response that eliminates dead cells and amyloid deposits and facilitates tissue repair.

### The coagulation, kallikrein/kinin and renin/angiotensin systems

Amyloid protofibrils have been shown to activate coagulation FXII, initiating contact system signaling and contributing to inflammation and thrombosis - particularly in Alzheimer’s disease.^50^ Although this contribution has not yet been confirmed in ATTR, the structural properties of transthyretin fibrils suggest the occurrence of a similar mechanism and (potentially) the promotion of vascular inflammation and thrombosis within the amyloid-laden myocardium. Indeed, FXI and fibrinogen were strongly enhanced in ATTR+ EVs. Microvascular damage and thrombosis might be further accentuated by enhanced vasoconstriction, due to the imbalance between the renin/angiotensin and kallikrein/kinin systems in ATTR+ EVs. Our results are in line with reports of the frequent thrombotic and hemorrhagic complications of amyloidosis.^51–55^

### Apolipoprotein E

The upregulation of apolipoprotein E is consistent with its known involvement in ATTR cardiomyopathy, where it binds to amyloid fibrils and influences their deposition and stability.^7,56^ Apolipoprotein E might also modulate the dynamics of TTR aggregation, although the underlying mechanisms have yet to be characterized.

### IL12 receptor β2 chain (IL12RB2) and CCL17

The observed downregulation of IL12RB2 (a key mediator of Th1 differentiation and interferon gamma signaling) and greater expression of CCL17 (a chemokine promoting T helper type 2 (Th2) cell recruitment) indicates a potential shift in immune polarization within the myocardial microenvironment of ATTR cardiomyopathy. This particular environment might constitute an effort to limit tissue damage and to facilitate wound healing, as seen in the canonical Th2 responses driven by helminth infections.^57^ However, Th2 responses are also known to stimulate fibrosis.

Overall, the protein content of EVs did not phenocopy findings in whole plasma, as exemplified by TTR. The differential abundance of proteins previously isolated from myocardial tissue in ATTR cardiomyopathy suggests that EVs can be released by cells activated or stressed by amyloid deposition, such as cardiomyocytes, endothelial cells and phagocytic cells. Thus, the EV content might represent a signature of local tissue remodeling and cellular dysfunction that is indicative of the disease stage.

## STUDY LIMITATIONS

Firstly, the small number of patients included in this pilot study precluded the establishment of a validation cohort and prevented a statistically robust comparison of ATTRv vs. ATTRwt patients; these two groups did not differ significantly in our analysis. Secondly, the cellular origin of the EVs (e.g., blood leukocytes, endothelial cells, or cells from amyloid-infiltrated tissues) could not be determined with precision. Thirdly, and although demographic, clinical, biomarker, and echocardiographic matching was carefully checked, differences between the groups persisted for certain variables: biomarkers (GGT and CRP), echocardiographic features (left ventricular ejection fraction (LVEF%)), and medications. These discrepancies might have introduced bias into group comparisons. Notably, the higher serum CRP concentrations observed in ATTR+ patients might have reflected a greater level of inflammation associated with amyloid deposition. This hypothesis is consistent with the elevated levels of complement proteins observed in ATTR+ EVs. Furthermore, the more frequent use of angiotensin-converting enzyme inhibitors (ACEIs), angiotensin II receptor blockers (ARBs), and mineralocorticoid receptor antagonists in ATTR- patients might have been a confounding factor in our analysis of the proteins from the renin-angiotensin and kallikrein-kinin systems. Notably, six ATTR+ patients had angiotensin/angiotensinogen levels exceeding 170 arbitrary units (AU), which accounted for most of the ATTR+ vs. ATTR- difference observed on the group level. In the remaining 35 ATTR+ patients, the mean ± standard deviation angiotensin/angiotensinogen level was 36.5 ± 19.4 AU. Four of the six individuals with elevated angiotensin/angiotensinogen levels were receiving ACE inhibitors or ARBs - a proportion similar to that observed in the ATTR+ group as a whole. However, all six exhibited severe biventricular dysfunction, which also affected liver function.

## CONCLUSION

Our study of plasma EVs from 41 patients with ATTR cardiomyopathy and 24 patients with non-amyloid hypertrophic cardiac disease led to the identification of a specific proteomic signature comprising 117 DEPs, most of which were upregulated in ATTR+ patients. Several of these proteins have been detected previously in amyloid-infiltrated tissues or are known to be involved in amyloidogenesis or the removal of amyloid fibrils. Notably, we discovered a tightly interconnected network centered on plasmin/plasminogen and involving the complement, coagulation, kallikrein-kinin, and renin-angiotensin systems. These systems might contribute to amyloid deposition, inflammation, and the cardiovascular risk.

In summary, the present findings may open up promising avenues for biomarker discovery and therapeutic targeting. Along with mechanistic investigations, validation in larger, independent cohorts is now required.

## CLINICAL PERSPECTIVES

### Clinical competencies in medical knowledge

Along with the identification of specific proteins of interest, the results of this study emphasize the value of EV-based proteomics for exploring disease-specific tissue processes in systemic conditions like ATTR cardiomyopathy and without the need for an invasive organ biopsy.

### Translational outlook

By identifying a network of proteins involved in the pathophysiology of ATTR heart disease through the analysis of plasma EVs alone, our work might foster the development of new tools for diagnosis and/or for disease stage assessment. Furthermore, some of the proteins identified here might be therapeutic targets. The detection of an elevated level of EV-associated TTR in ATTR+ patients (despite the presence of low plasma TTR levels) further underscores the diagnostic potential of EV analysis.

## ACKNOWLEDGEMENTS

We sincerely thank the Amyloidosis Foundation and the Amyloidosis Network for their invaluable support and contributions to this research. We are also grateful to Région Île-de-France Cancéropôle and Université Paris-Est-Créteil for their contributions to the acquisition of the nano-LC-MS platform (NanoAcquity-ESI-SynaptG2-Si, Waters, Milford, MA, USA).

## ABBREVIATIONS

α1AT: alpha-1 antitrypsin
α2AP: alpha-2 antiplasmin
ACEI: angiotensin-converting enzyme inhibitor
AGT: angiotensinogen
ALT: alanine aminotransferase
ARB: angiotensin II receptor blocker
AST: aspartate aminotransferase
ATTR: TTR amyloidosis
ATTRv: TTR amyloidosis, variant form
ATTRwt: TTR amyloidosis, wild-type form
AU: arbitrary unit
bpm: beats per minute
CFB: complement factor B
CFD: complement factor
D CRP: C-reactive protein
DEP: differentially expressed protein
DOID: Disease Ontology Identifier
ECM: extracellular matrix
eGFR: estimated glomerular filtration rate
EV: extracellular vesicle
FDR: false discovery rate
GO: Gene Ontology
GGT: gamma-glutamyl transferase
GLS: global longitudinal strain
HR: heart rate
IQR: interquartile range
IVS: interventricular septum
LC-MS: liquid chromatography mass spectrometry
LVEF: left ventricular ejection fraction
NT-proBNP: N-terminal pro-brain natriuretic peptide
NYHA: New York Heart Association
PASP: pulmonary artery systolic pressure
PLG: plasmin/plasminogen
PTX3: pentraxin 3
SBP: systolic blood pressure
SERPINF2: alpha-2-antiplasmin
TTR: transthyretin

---

**Figure S1.**
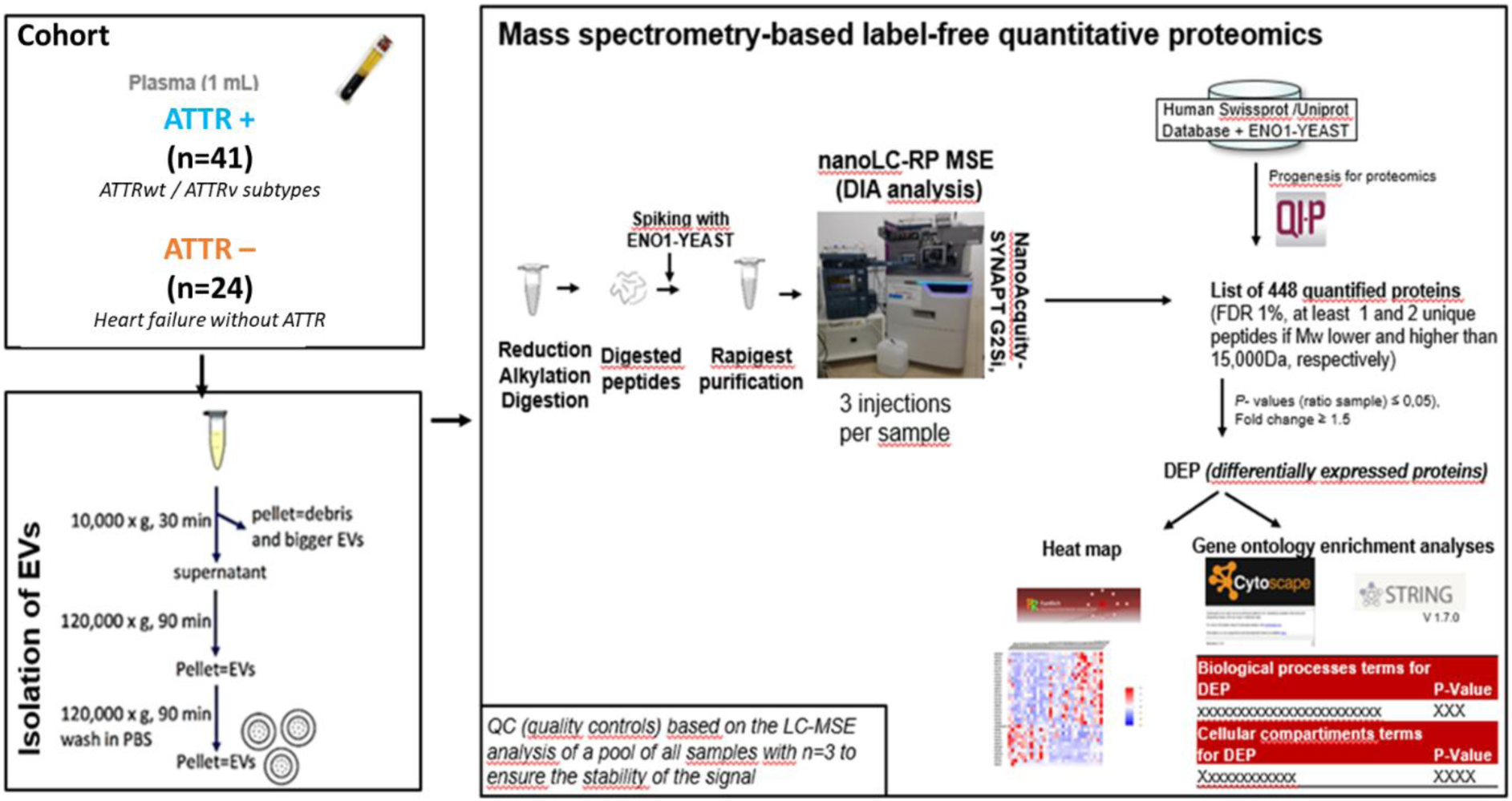
The shotgun proteomics workflow for characterizing the circulating extracellular protein content in amyloidosis.

**Figure S2.**
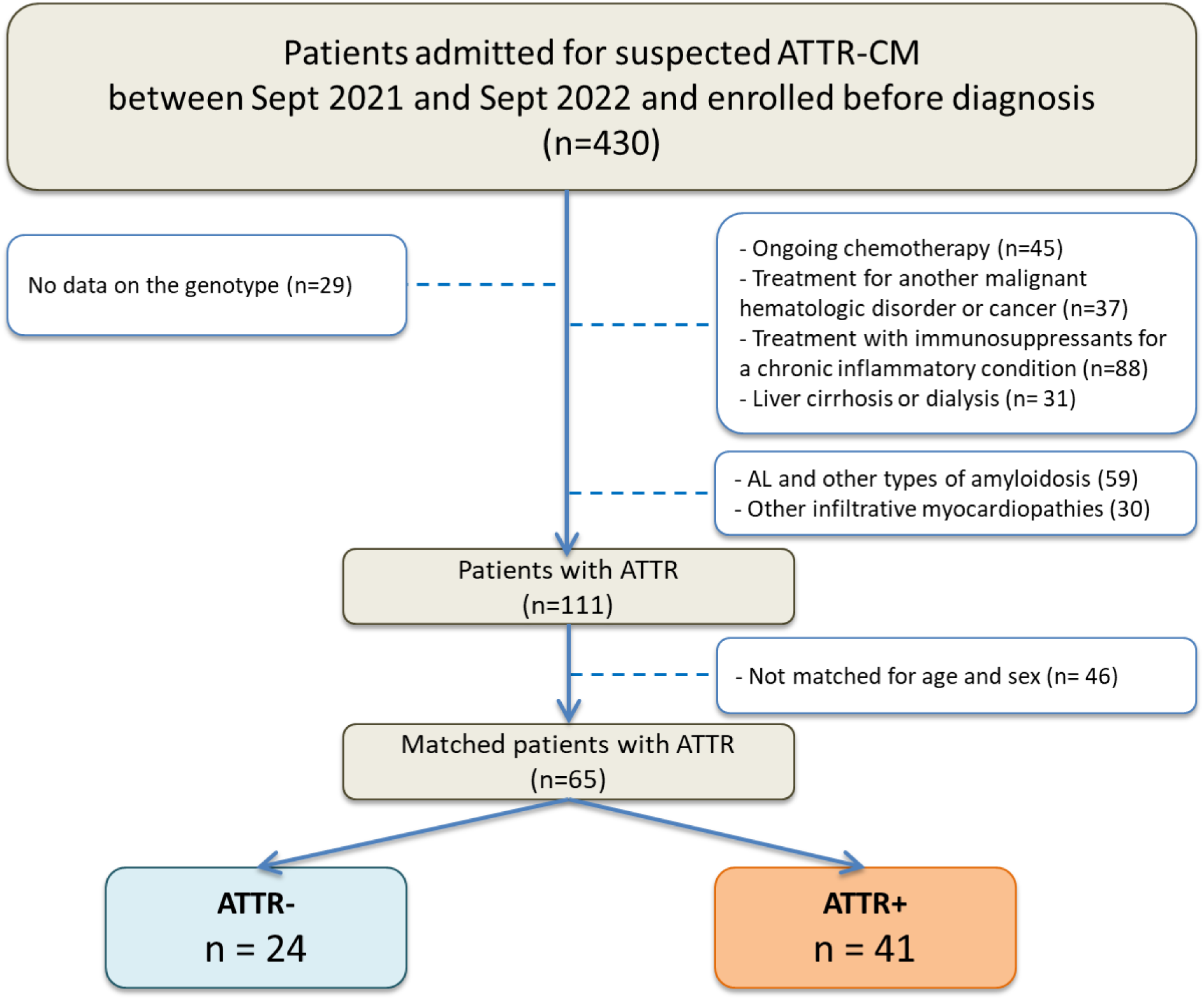
Study flowchart.

**Figure S3.**
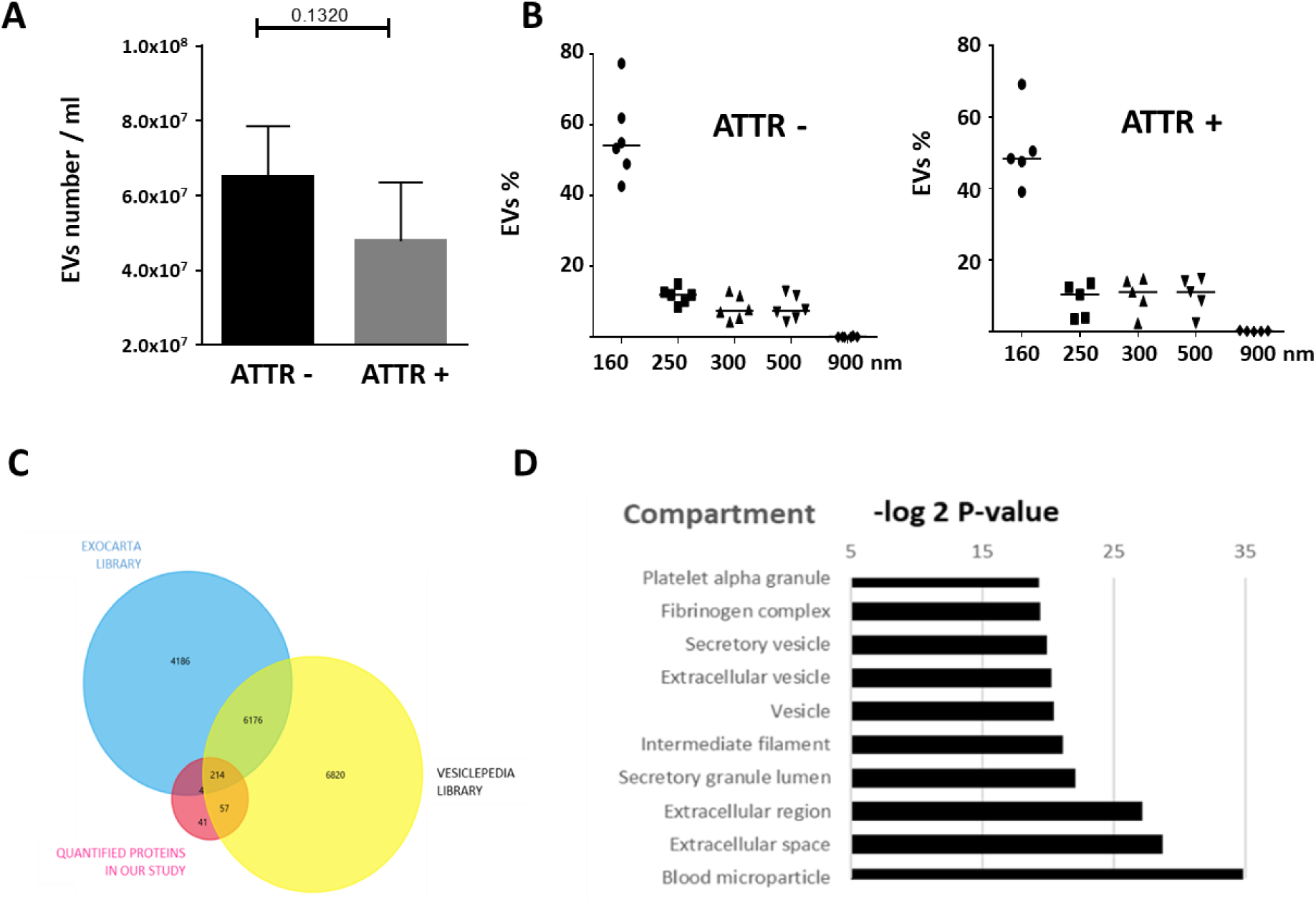
Characteristics of extracellular vesicles (EVs) derived from plasma samples from ATTR+ and ATTR- patients. (A) The number of EVs in plasma was assessed by flow cytometry, using Trucount^TM^ beads. EVs were first identified by CD9 labeling. B) EV size in the **ATTR+ and ATTR- groups.** Megamix fluorescent beads were used to differentiate between particles with diameters of 160, 200/240, 300, 500 and 900 nm. The flow cytometry data were analyzed with FlowJo software (FlowJo, Ashland, OR; version 10.7.1). (C) A Venn diagram of the quantified proteins found in the Vesiclepedia and/or ExoCarta EV libraries. (D) The top 10 cellular/extracellular compartments in the GO enrichment analysis, using StringApp (version 2.1.1) implemented in Cytoscape (version 3.9.1).

**Figure S4.**
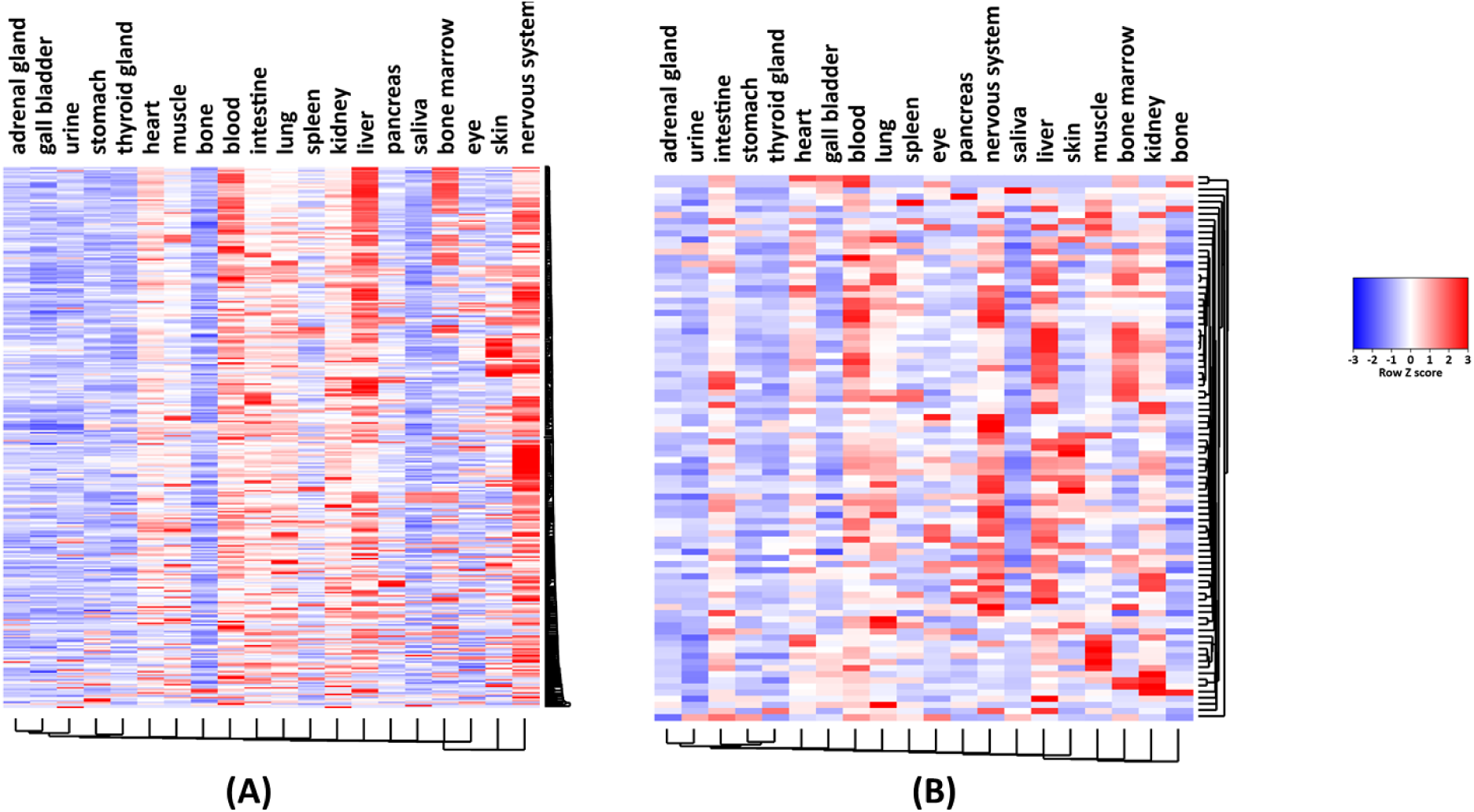
Heat maps representing the tissue/blood cell origins of EVs in plasma samples for (A) all quantified proteins (337 of the 448 proteins mapped in StringApp, version v2.1.1). (B) EV proteins in the ATTR signature (89 of the 117 DEPs mapped in StringApp ). Heat maps were drawn with Funrich software, using String data. The Z-score was calculated from the String confidence score for protein localization in the Compartments database (https://compartments.jensenlab.org/Search). A higher score means a higher degree of confidence.

**Figure S5.**
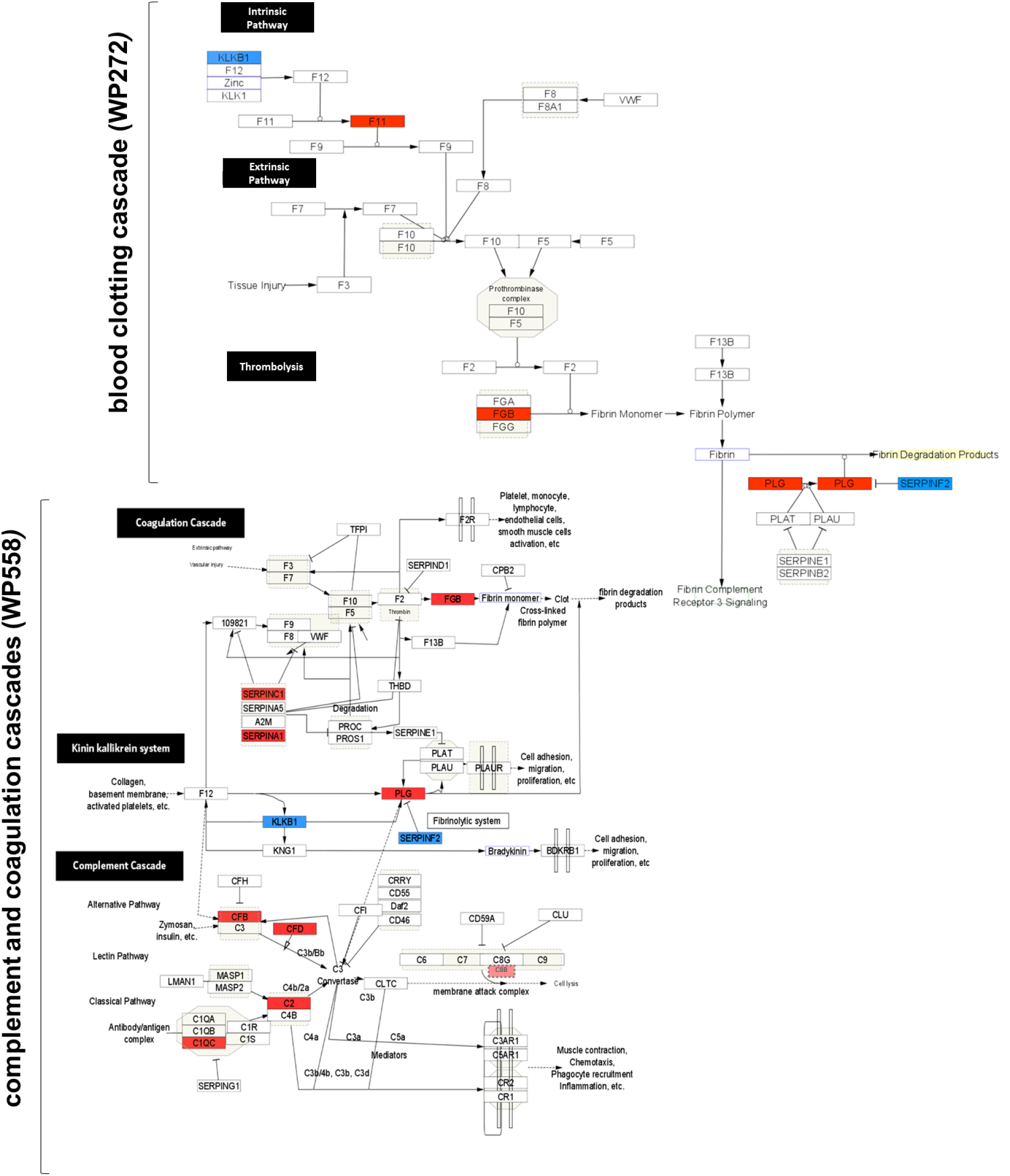
Proteins deregulated in ATTR+ EVs in the complement and coagulation cascades (WP558 - Homo sapiens - version 2024-07-22) and blood clotting cascade (WP272 - Homo sapiens - version 2023-11-25; wikipathway: https://www.wikipathways.org/pathways). Proteins upregulated in ATTR+ EVs (relative to ATTR- EVs) are highlighted in red, while proteins downregulated in ATTR+ EVs (relative to ATTR- EVs) are highlighted in blue.

