## Supplementary material for "QUANTITATIVE PROTEOMICS OF PLASMA EXTRACELLULAR VESICLES REVEALS A TTR-PLASMINOGEN NETWORK IN ATTR CARDIAC AMYLOIDOSIS": supp data 1

**SUPPLEMENTAL DATA**

### Supplemental data 1: Detailed material and methods for EV proteomics analysis

#### 1. Extracellular vesicle (EV) isolation

The protocol is based on the isolation of EVs prior to characterization by FACS and proteomics, as described previously with some modifications^1^ and in line with the MISEV guidelines.^2,3^ Briefly, 1 mL of plasma was centrifuged at 200 g for 5 min and then at 2,000 g for 10 min, to remove cell debris. Supernatant fractions were centrifuged at 12,000 g for 30 min. EVs were obtained from the supernatant by centrifugation at 120,000g for 90 min (Beckman Ti50). The resulting pellets were resuspended in a large volume of ultrapure water filtered through a 220-nm filter, and then washed and collected by ultracentrifugation at 120,000 g for 90 min. All the steps were performed at 4°C. EV-containing pellets were resuspended in 200 μL of ultrapure water and kept overnight at 4°C. The concentration of EV proteins was determined using a micro-BCA Protein Assay Kit (Thermo Fisher Scientific, Waltham, MA, USA). The EV preparations were stored at -80°C.

#### 2. Flow cytometry analysis of EVs

Fluorescence-based EV counts in plasma were determined on an LSR Fortessa ﬂow cytometer (BD Biosciences, Le Pont de Claix, France), using Trucount beads. EVs were preliminarily identified by CD9 labeling. For the size differentiation of EVs, Megamix ﬂuorescent beads were used for particles with diameters of 160, 200/240, 300, 500 and 900 nm. The ﬂow cytometry data were analyzed with FlowJo software (version 10.7.1, FlowJo, Ashland, OR, USA).

All statistical analyses were performed with Prism 6.07 software (GraphPad Software, La Jolla, CA, USA). Differences were evaluated in a Mann-Whitney test.

#### 3. Label-free quantitative proteomic analysis

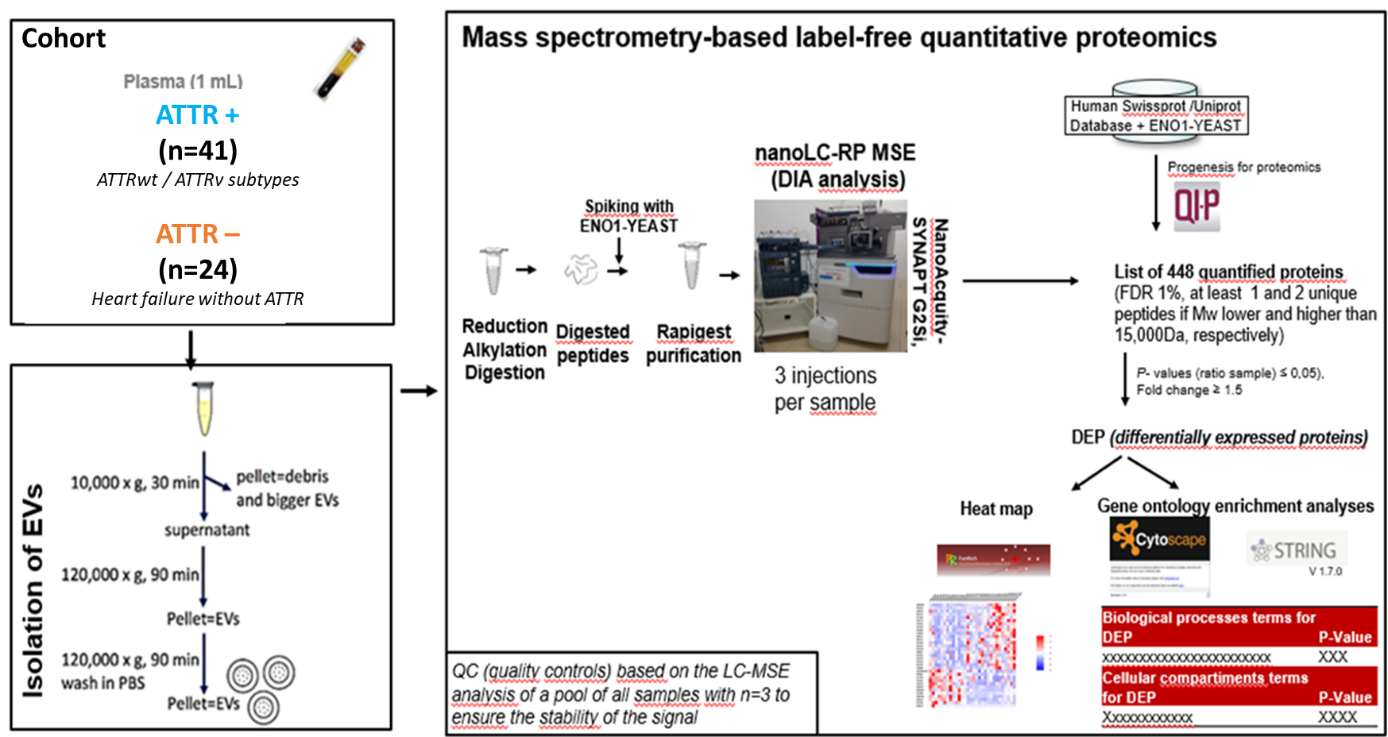

Figure S1. Shotgun proteomics workflow for characterizing the circulating extracellular protein content in amyloidosis.

A differential, label-free, quantitative mass spectrometry (MS) analysis was performed as follows. First, 30 μg of isolated EVs were diluted in 50 mM ammonium bicarbonate pH 7.8 with Rapigest (Waters, Milford, MA, USA). Next, reduction, alkylation and digestion were performed with dithiothreitol, iodoacetamide, and trypsin, respectively. Tryptic digestion was performed overnight at 37°C with a trypsin/protein ratio of 3%. Formic acid (FA) was added to each sample (final concentration: 0.3%) to stop the trypsin digestion and obtain pellets. The supernatant (containing desalted peptides) was retrieved and dried, using a Speed-Vac. Dried samples were diluted in buffer A (H_2_O with 0.1% FA) and analyzed using nano-LC-MS on a NanoAcquity-ESI-SynaptG2-Si system (Waters) operating in nano-ESI positive mode. After loading onto the UPLC peptide BEH130 C18 nanoACQUITY^TM^ column (100 μm x 100 mm, diameter: 1.7 μm diameter; Waters) at a flow rate of 0.45 μL/min, peptides were separated with a gradient increasing from 1% Buffer B (acetonitrile with 0.1% FA)/99% Buffer A (H_2_O with 0.1% FA) to 40% Buffer B/60% Buffer A, and then to 85% Buffer B/15% Buffer A, over 120 min. MS spectra were recorded in a 50-2,000 m/z mass range at a resolution of 10,000-20,000. The nano LC-MS method included 120 min of acquisition time, positive polarity, MS acquisition range of 50-4,000 Da, and a collision energy range of 20-55 V. Proteins were identified and quantified using Progenesis software for proteomics QI (Waters), with the ion accounting algorithm and the following parameters: trypsin digestion enzyme; a maximum of two missed cleavages; maximum protein mass: 250 kDa; fixed modification: carbamodomethyl (C); variable modifications: oxidation (M), deamination (N, Q) and acetylation (N-ter); false discovery rate: 1%, automatic mass tolerance, isotope filter: 2; *Homo sapiens* UniprotKB/Swissprot database (canonical proteins from UP000005640 proteome Release-2021-02; https://www.uniprot.org/) concatenated with the sequence of angiotensin II (DRVYIHPF) and the internal standard (ENO1_YEAST, P00924). The proteomic data were A analyzed quantitatively in an analysis of variance , using the Progenesis for proteomics QI application (Waters). We focusing on the three most abundant and confidently identified unique peptides (non-conflicting peptides) by considering the correlation factor of each and normalizing the signal against the ENO-1 internal standard. Differential quantitative data were visualized in a volcano plot, in order to determine and justify the fold change (FC) threshold for significant protein deregulations in each corresponding dataset (Table S1). Therefore, only proteins with p≤0.05 and a FC>1.25 were considered to be significantly deregulated. These differentially expressed proteins were included in the EV-ATTR signature.

#### 4. Downstream analysis of proteomic data

To check the proportion of proteins known to be found in EVs, the experimental data were compared with EV libraries (ExoCarta, July_2015_Release ^4–6^ and/or Vesiclepedia, August_2018_Release ^7,8^. These results were plotted as Venn diagrams, using FunRich software (version 3.1.3) ^9,10^.

To reduce the dimensionality of the dataset for each quantified protein from the 65 samples to two dimensions and determine whether or not the protein profiles from ATTR+ and ATTR- patients forms two distinct clusters, we performed a principal component analysis on the EV-ATTR signature using R (version 4.2.2) with the ggfortify and ggplot2 packages and the “fviz_pca_ind” function. The heat map was plotted using FunRich software (version 3.1.3) ^9,10^.

To generate both EV-ATTR network and GO/pathway enrichment analyses, we used StringApp v2.1.1 ^11^ implemented in Cytoscape v3.9.1 ^12^. The STRING network of the EV-ATTR signature was clustered through the MCL cluster network, with a granularity parameter of 4. A gene ontology (GO) enrichment analysis was performed with *Homo sapiens* database as the background (Table S2). The GO cellular compartment enrichment analysis was performed for all the identified proteins, while the GO biological processes and pathways enrichment analyses were performed for the EV-ATTR signature STRING network only (i.e. significantly deregulated proteins in ATTR *vs.* non-amyloid cardiomyopathy).
