## Supplementary figures and images for "QUANTITATIVE PROTEOMICS OF PLASMA EXTRACELLULAR VESICLES REVEALS A TTR-PLASMINOGEN NETWORK IN ATTR CARDIAC AMYLOIDOSIS"

### supp data 2

#### Supplemental data 2: STUDY Flowchart

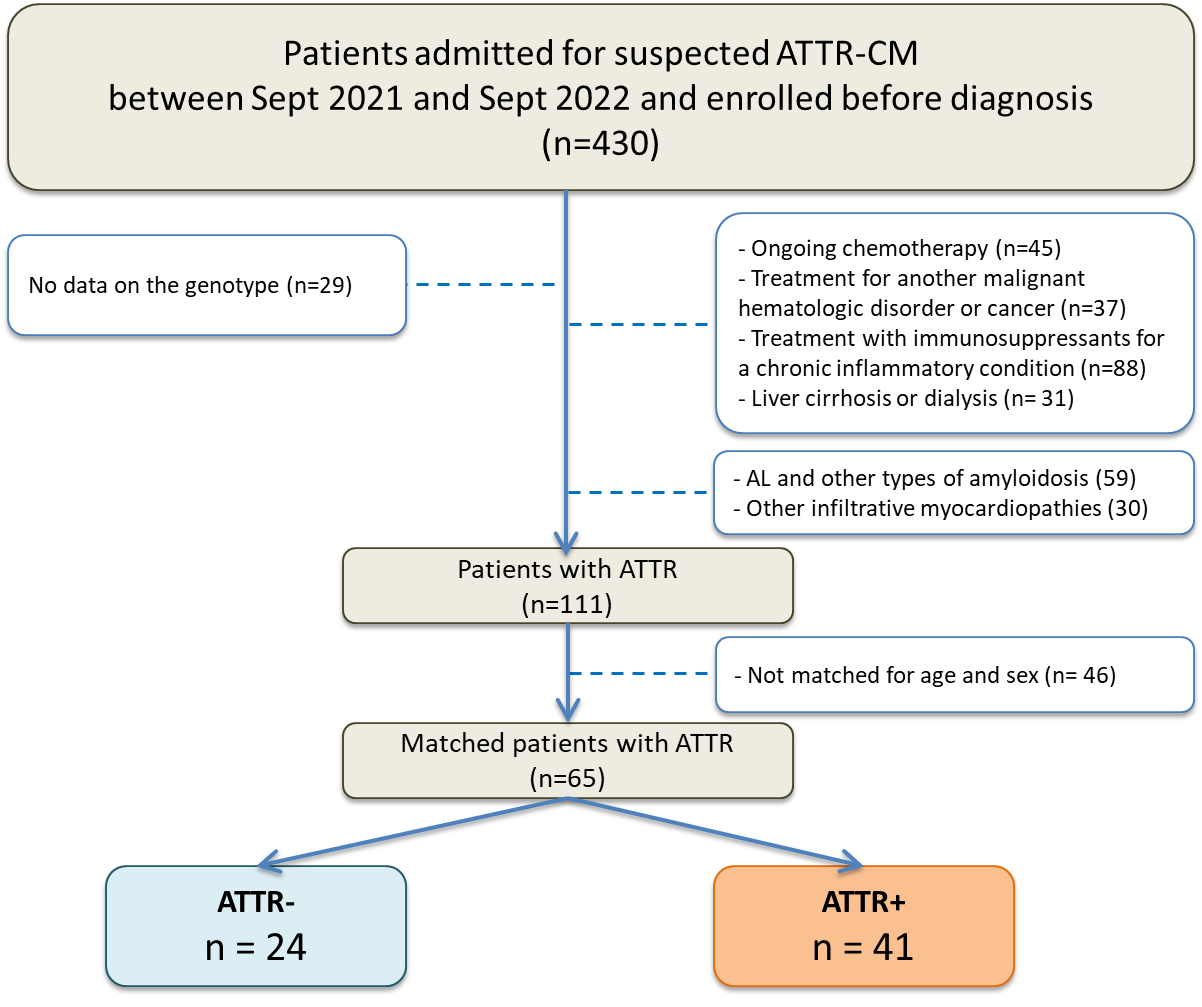
