## Supplementary material for "QUANTITATIVE PROTEOMICS OF PLASMA EXTRACELLULAR VESICLES REVEALS A TTR-PLASMINOGEN NETWORK IN ATTR CARDIAC AMYLOIDOSIS": supp data 3

### Supplemental data 3: EV charaCteriZation and purity

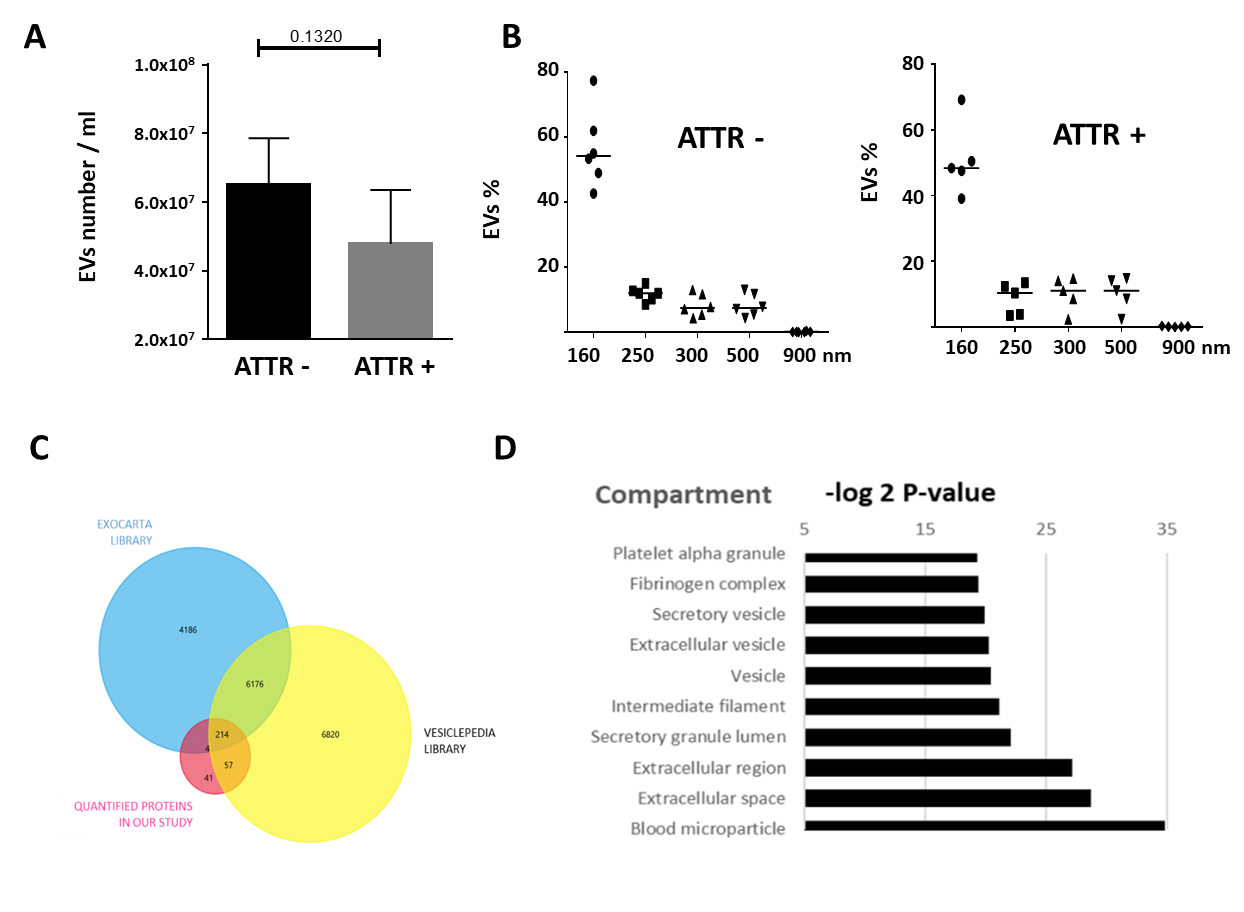
 **Figure S3. Characterization of EVs derived from plasma from ATTR+ and ATTR- patients.** (A) The number of EVs in plasma was assessed by flow cytometry, using Trucount beads. The EVs were preliminarily identified by CD9 labeling. B) For the size differentiation of EVs, Megamix ﬂuorescent beads were used for particles with diameters of 160, 200/240, 300, 500 and 900 nm. The ﬂow cytometry data were analyzed with FlowJo software (version 10.7.1, FlowJo, Ashland, OR). (C) A Venn diagram representing the quantified proteins found in the Vesiclepedia^2,3^ and/or ExoCarta^4^ EV libraries (D) The top 10 extracellular and cellular compartments in the GO enrichment analysis, using StringApp v2.1.1 implemented in Cytoscape v3.9.1.

The EV concentration and EV size distribution were similar in the ATTR+ and ATTR groups. In both groups, most of the EVs were 160 nm in diameter, indicating that they were primarily exosomes (small cell microparticles typically derived from intracellular compartments), rather than larger cell microparticles typically associated with the plasma membrane.^1^

The Venn diagram shows that 87% of the mapped, quantified proteins had already reported in Vesiclepedia^2,3^ and/or ExoCarta^4^ EV libraries. Overall, this analysis confirmed the high purity of EVs. A GO enrichment analysis of the extracellular/cellular compartments confirmed that the majority of the proteins were related to EVs. Ranked by significance, the top 10 extracellular/cellular compartments (all with a p-value below 0.00001) were as follows: extracellular space, blood microparticle, extracellular region, secretory granule lumen, vesicle, EV, extracellular membrane-bounded organelle, secretory granule, fibrinogen complex, and platelet alpha granule.
