## Supplementary material for "QUANTITATIVE PROTEOMICS OF PLASMA EXTRACELLULAR VESICLES REVEALS A TTR-PLASMINOGEN NETWORK IN ATTR CARDIAC AMYLOIDOSIS": supp data 4

### Supplemental data 4. Distribution enrichment analysis of THE EV-ATTR Protein Signature IN Tissues and Body Fluids.

Circulating EVs represent a complex and heterogeneous system, as they can be derived from any cell or tissue in the body. We tried to characterize the source of the plasma EVs by mapping the tissue and blood distribution of total proteins and the differentially expressed proteins (DEPs) (Figure S4A and B, respectively). Most of the proteins were present in the blood, the heart, and several other organs commonly infiltrated by amyloid fibrils, such as the peripheral nervous system, liver, and kidney.^1^ Thus, plasma EV proteins might reflect the protein content of the main target organs in the amyloidogenic process.

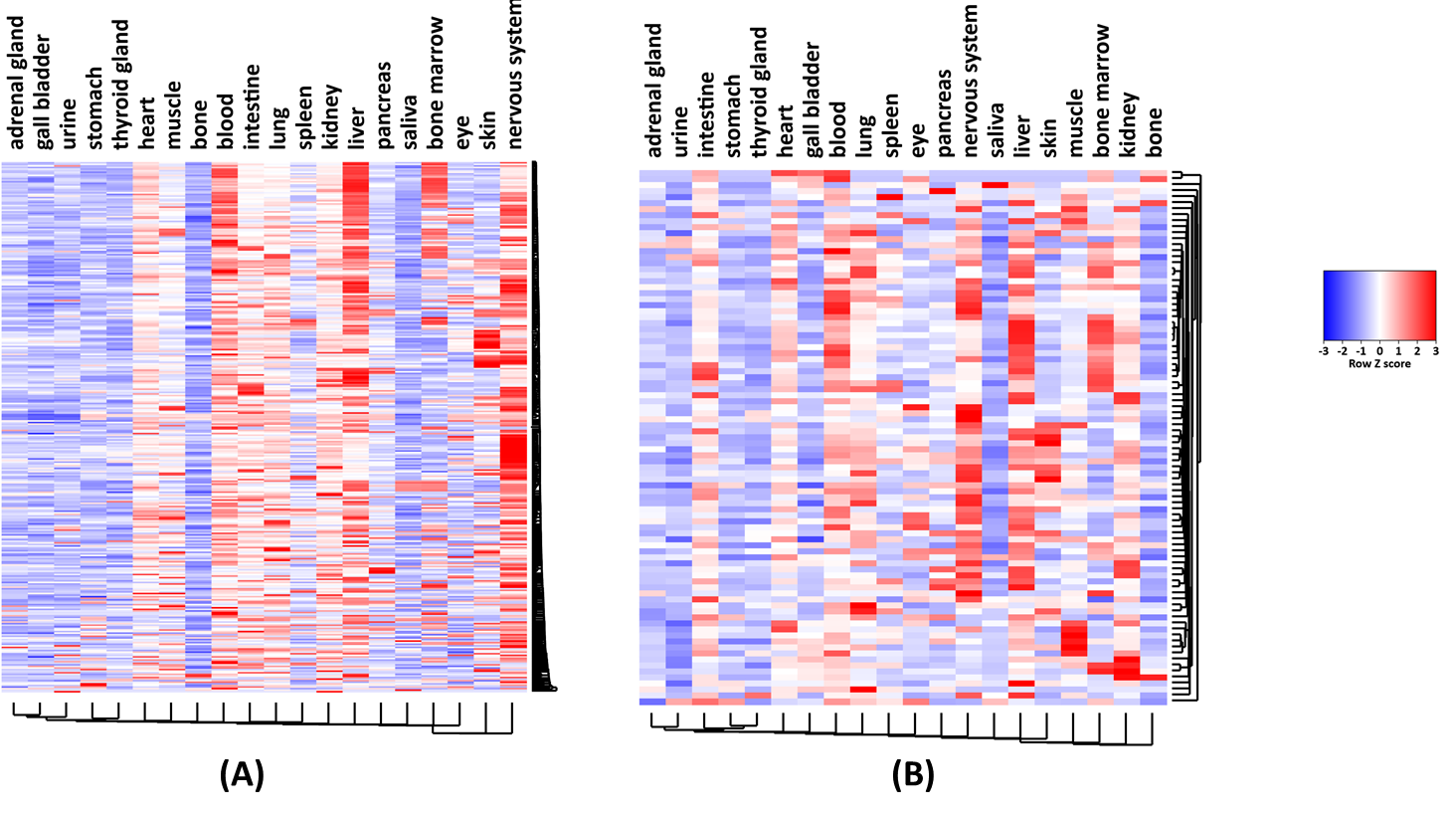

Fig S4. Heat maps representing the landscape of tissue/blood cell origins among plasma EV samples for (A) all quantified proteins (337 of the 448 proteins mapped in STRING), (B) EV proteins in the ATTR signature (89 of the 117 of DEPs mapped in STRING). The heat maps were plotted using Funrich software and STRING data (StringApp v2.1.1). The Z-score was calculated from the STRING confidence score from the Compartments database (https://compartments.jensenlab.org/search); a higher score means a higher degree of confidence.)
