## Supplementary material for "QUANTITATIVE PROTEOMICS OF PLASMA EXTRACELLULAR VESICLES REVEALS A TTR-PLASMINOGEN NETWORK IN ATTR CARDIAC AMYLOIDOSIS": supp data 5

### Slide 1
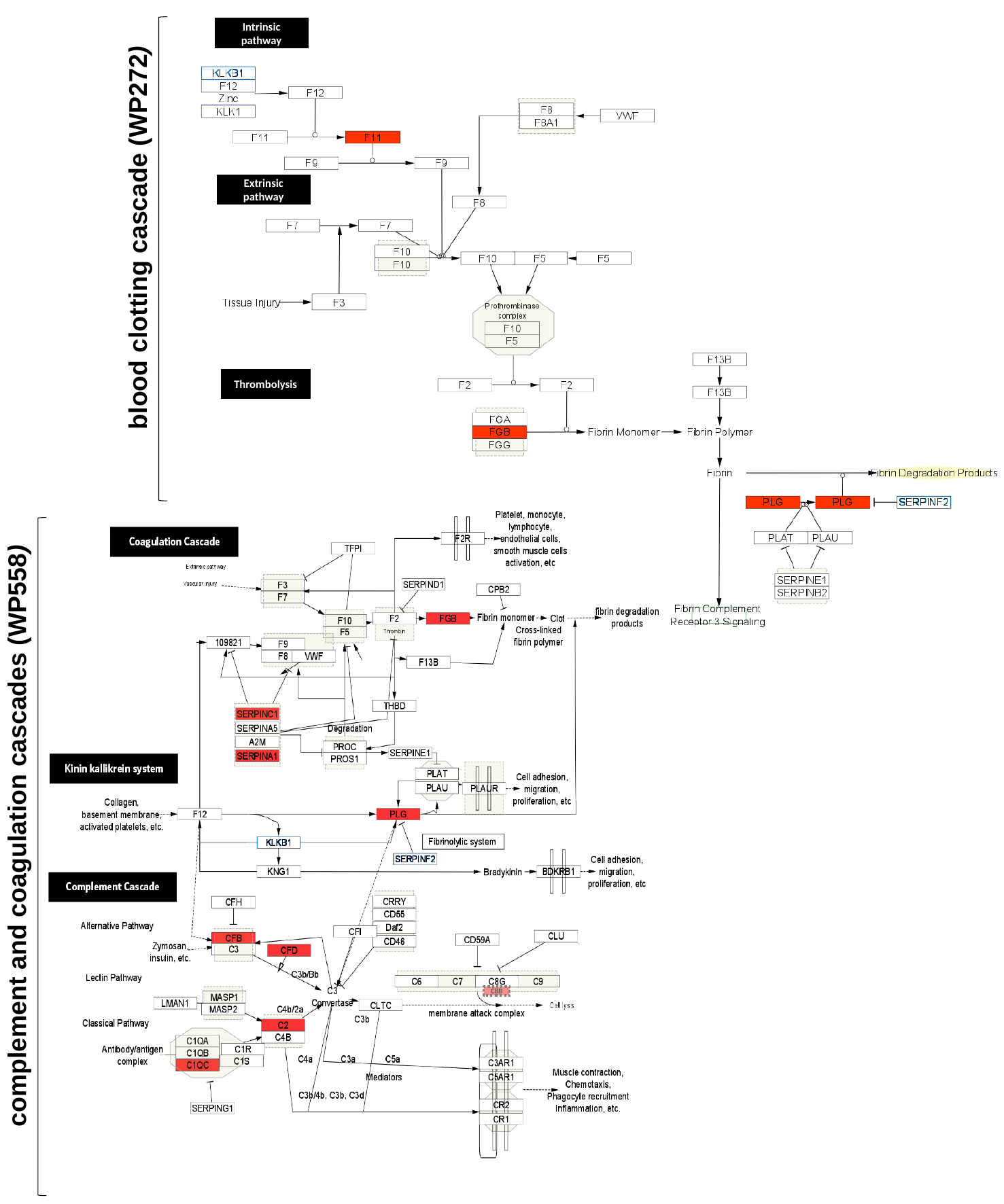

Intrinsic pathway
Extrinsic pathway
blood clotting cascade (WP272)
Thrombolysis
C8B
complement and coagulation cascades (WP558)
